# Experimental re-infected cats do not transmit SARS-CoV-2

**DOI:** 10.1101/2021.01.18.427182

**Authors:** Natasha N. Gaudreault, Mariano Carossino, Igor Morozov, Jessie D. Trujillo, David A. Meekins, Daniel W. Madden, Konner Cool, Bianca Libanori Artiaga, Chester McDowell, Dashzeveg Bold, Velmurugan Balaraman, Taeyong Kwon, Wenjun Ma, Jamie Henningson, Dennis W. Wilson, William C. Wilson, Udeni B. R. Balasuriya, Adolfo García-Sastre, Juergen A. Richt

## Abstract

SARS-CoV-2 is the causative agent of COVID-19 and responsible for the current global pandemic. We and others have previously demonstrated that cats are susceptible to SARS-CoV-2 infection and can efficiently transmit the virus to naïve cats. Here, we address whether cats previously exposed to SARS-CoV-2 can be re-infected with SARS-CoV-2. In two independent studies, SARS-CoV-2-infected cats were re-challenged with SARS-CoV-2 at 21 days post primary challenge (DPC) and necropsies performed at 4, 7 and 14 days post-secondary challenge (DP2C). Sentinels were co-mingled with the re-challenged cats at 1 DP2C. Clinical signs were recorded, and nasal, oropharyngeal, and rectal swabs, blood, and serum were collected and tissues examined for histologic lesions. Viral RNA was transiently shed via the nasal, oropharyngeal and rectal cavities of the re-challenged cats. Viral RNA was detected in various tissues of re-challenged cats euthanized at 4 DP2C, mainly in the upper respiratory tract and lymphoid tissues, but less frequently and at lower levels in the lower respiratory tract when compared to primary SARS-CoV-2 challenged cats at 4 DPC. Histologic lesions that characterized primary SARS-CoV-2 infected cats at 4 DPC were absent in the re-challenged cats. Naïve sentinels co-housed with the re-challenged cats did not shed virus or seroconvert. Together, our results indicate that cats previously infected with SARS-CoV-2 can be experimentally re-infected with SARS-CoV-2; however, the levels of virus shed was insufficient for transmission to co-housed naïve sentinels. We conclude that SARS-CoV-2 infection in cats induces immune responses that provide partial, non-sterilizing immune protection against reinfection.

## Introduction

Severe Acute Respiratory Syndrome Coronavirus 2 (SARS-Cov-2) is the causative agent of Coronavirus Disease 2019 (COVD-19), first identified in Wuhan China in late 2019 and responsible for the ongoing global pandemic (1). SARS-CoV-2 is highly transmissible and capable of causing severe disease in humans. Furthermore, there have been multiple cases reported of transmission from COVID-19 patients to animals including cats, large cats, dogs, ferrets, and mink in China, South America, the United States and Europe (2-9, summary of US cases: https://www.aphis.usda.gov/aphis/ourfocus/animalhealth/sa_one_health/sars-cov-2-animals-us). Evidence supporting reverse zoonosis of mink infecting humans has also been reported (10). Understanding SARS-CoV-2 susceptibility, transmission and re-infection in companion animals and livestock species that are frequently in close proximity with humans is important for assessing risk and implementing mitigation strategies to stop virus spread in order to maintain public health as well as food and economic security (11).

Recently, we and others have demonstrated that domestic cats are susceptible to SARS-CoV-2 by experimental infection and can readily transmit the virus to naïve cats (12-15). Cats inoculated via natural routes of exposure are acutely infected and shed the virus from nasal, oral and rectal cavities starting from 1 up to 14 days with peak virus shedding occurring within the first 7 days after infection (12-15). Together these studies show that cats ranging from 4 months up to 8 years of age remain asymptomatic with no significant gross pathological changes, and mild to moderate histological alterations associated mainly with the upper respiratory tract tissues (12-15). In contrast, mortality and severe histological lesions in tissues of the upper and lower respiratory tract were observed for juvenile cats younger than 4 months infected with SARS-CoV-2 (13). Cats also develop virus-specific and neutralizing antibody responses to SARS-CoV-2 (7, 12-17). A more detailed understanding of the role that these immune responses play in protection from re-exposure to SARS-CoV-2 is imperative.

Here, we present a detailed investigation on SARS-CoV-2 re-infection of sub-adult domestic cats at 21 DPC in two independent studies. In the second re-infection study, 2 sentinel contact cats were introduced at 1 DP2C to determine if transmission to naïve animals could occur after re-infection. Results of the clinical course of SARS-CoV-2 re-infection, viral shedding and transmission cats previously infected with SARS-CoV-2 are presented.

## Materials and Methods

### Cells and Virus

Vero E6 cells (ATCC^®^ CRL-1586^™^, American Type Culture Collection, Manassas, VA, USA) were used for virus propagation and titration. Cells were cultured in Dulbecco’s Modified Eagle’s Medium (DMEM, Corning, New York, N.Y, USA), supplemented with 5% fetal bovine serum (FBS, R&D Systems, Minneapolis, MN, USA) and antibiotics/antimycotics (ThermoFisher Scientific, Waltham, MA, USA), and maintained at 37 °C under a 5% CO_2_ atmosphere. The SARS-CoV-2 USA-WA1/2020 strain was acquired from BEI Resources (Manassas, VA, USA) and passaged 3 times in Vero E6 cells to establish a stock virus (1×10^6^ TCID_50_/ml) for inoculation of animals. This stock virus was sequenced by next generation sequencing (NGS) using the Illumina MiSeq and its consensus sequence was found to be homologous to the original USA-WA1/2020 strain (GenBank accession: MN985325.1). To determine infectious virus titer, 10-fold serial dilutions were performed on Vero E6 cells. The presence of cytopathic effects (CPE) after 96 hours incubation was used to calculate the 50% tissue culture infective dose (TCID_50_)/ml using the Spearman-Karber method.

### Animals and experimental design

#### Ethics statement for use of animals

All animal studies and experiments were approved and performed under the Kansas State University (KSU) Institutional Biosafety Committee (IBC, Protocol #1460) and the Institutional Animal Care and Use Committee (IACUC, Protocol #4390) in compliance with the Animal Welfare Act. All animal and laboratory work were performed in biosafety level-3+ and -3Ag laboratory and facilities in the Biosecurity Research Institute at KSU in Manhattan, KS, USA.

#### SARS-CoV-2 re-infection of animals

Two independent re-infection studies were performed in sub-adult domestic cats 5 or 7 months of age, respectively. Animal identification numbers and treatment assignments for each study are summarized in **Table 1**. In the first study, 3 cats from a previous SARS-CoV-2 challenge and transmission study (12), including 1 principal and 2 sentinel cats, were re-challenged with SARS-CoV-2 at 21 days post primary challenge (DPC) and sacrificed at 4 days post second challenge (DP2C). The same dose as primary challenge of 2ml of 10^6^ TCID_50_ SARS-CoV-2 was administered to each cat both intranasal (0.5ml per nostril) and per oral (1ml) routes.

**Table 1.** Animals and treatment assignments.

| **Study 1** (5-month-old domestic cats) |  |  |  |  |
| --- | --- | --- | --- | --- |
| Primary challenge |  |  | Re-challenge |  |
| *Cat ID* | *Treatment* | *Necropsy* | *Treatment* | *Necropsy* |
| 328 | principal | na | principal (21 DPC) | 4 DP2C |
| 272 | sentinel (1 DPC) | na | principal (21 DPC) | 4 DP2C |
| 903 | sentinel (1 DPC) | na | principal (21 DPC) | 4 DP2C |
| **Study 2** (7-month-old domestic cats) |  |  |  |  |
| Primary challenge |  |  | Re-challenge |  |
| *Cat ID* | *Treatment* | *Necropsy* | *Treatment* | *Necropsy* |
| 305 | principal | 4 DPC | na | na |
| 526 | principal | 4 DPC | na | na |
| 160 | principal | 21 DPC | na | na |
| 135 | principal | na | principal (21 DPC) | 4 DP2C |
| 590 | principal | na | principal (21 DPC) | 4 DP2C |
| 247 | principal | na | principal (21 DPC) | 7 DP2C |
| 119 | sentinel (1 DPC) | na | principal (21 DPC) | 7 DP2C |
| 127 | sentinel (1 DPC) | na | principal (21 DPC) | 14 DP2C |
| 280 | na | na | sentinel (1 DP2C) | 14 DP2C |
| 620 | na | na | sentinel (1 DP2C) | 14 DP2C |

principal=virus inoculated cat; sentinel=naïve contact cat; DPC=days post primary challenge; DP2C=days post second challenge; na=not applicable

In the second study, a total of 6 principal cats, 2 groups of 3, were challenged with SARS-CoV-2 at the same dose and route as detailed in the first study. At 1 DPC, 2 sentinel contact cats were added, 1 per group of challenged cats. Postmortem examinations were performed on two principal animals at 4 DPC and one at 21 DPC. At 21 DPC, 5 cats including the remaining 3 principals and the 2 sentinels were re-challenged with SARS-CoV-2 using same dose and route as in the primary challenge. At 1 DP2C, 2 new naïve sentinel cats were introduced. Two re-challenged animals each were euthanized and postmortem examinations performed at 4 and 7 DP2C, respectively. The remaining re-challenged cat and the 2 sentinels were euthanized and postmortem examinations performed at 14 DP2C.

### Clinical evaluations and sample collection

Cats were observed daily for clinical signs, as described previously (12). Weights of all cats were recorded on bleed days. Blood and serum were collected from all primary inoculated and sentinel cats at 0, 1, 3, 4, 7, 14 and 21 DPC, and the re-infected and sentinels at 0, 1, 3, 4, 7, 11 and 14 DP2C via venipuncture of the cephalic vein under anesthesia or during terminal bleeding by cardiac puncture. Nasal, oropharyngeal and rectal swabs were collected at 0, 1, 3, 4, 5, 7, 10, 14, 18 and 21 DPC, and at 0, 1, 3, 4, 5, 7, 8, 11 and 14 DP2C in 2 ml of DMEM (Corning,) with antibiotics/antimycotic (ThermoFisher). Swabs were vortexed and supernatant aliquoted directly into cryovials or into RLT buffer (Qiagen, Germantown, MD, USA) and stored at −80 ^°^C until further analysis.

A full postmortem examination was performed for each cat and gross changes (if any) were recorded. Tissues were collected either in 10% neutral-buffered formalin (Fisher Scientific, Waltham, MA, USA), or as fresh tissues which were then frozen at −80^°^C. Tissues were collected from the upper respiratory tract (URT) and lower respiratory tract (LRT), central nervous system (brain and cerebral spinal fluid [CSF]), gastrointestinal tract (GIT) as well as accessory organs. The lungs were removed *in toto* including the trachea, and the main bronchi were collected at the level of the bifurcation and at the entry point into the lung lobe. Lung lobes were evaluated based on gross pathology and collected and sampled separately. Bronchoalveolar lavage fluid (BALF), nasal wash and urine were also collected during postmortem examination and stored at −80^°^C until analyzed. Fresh frozen tissue homogenates were prepared as described previously (12) and the supernatant retained for RNA extraction and quantitative reverse transcription real-time PCR (RT-qPCR).

#### RNA extraction and quantitative real-time reverse transcription PCR (RT-qPCR)

SARS-CoV-2-specific RNA was detected using a quantitative reverse transcription real time -PCR (RT-qPCR) assay as previously described (12). Briefly, tissue homogenates in DMEM, blood, CSF, BALF, urine, and swab samples in DMEM were mixed with an equal volume of RLT RNA stabilization/lysis buffer (Qiagen, Germantown, MD, USA), and 200μl of sample lysate was then used for extraction using a magnetic bead-based nucleic acid extraction kit (GeneReach USA, Lexington, MA) on an automated Taco^™^ mini nucleic acid extraction system (GeneReach) as described previously (12). Positive (IDT, IA, USA; 2019-nCoV_N_Positive Control, diluted 1:100 in RLT) and negative extraction controls were employed.

Quantification of SARS-CoV-2 RNA was performed as previously described (12) using the N2 SARS-CoV-2 primer and probe set in a RT-qPCR protocol established by the CDC for the detection of SARS-CoV-2 nucleoprotein (N)-specific RNA. A 10-point standard curve of quantitated viral RNA (USA-WA1/2020 isolate) was used to quantify RNA copy number. A positive Ct cut-off of 40 cycles was used and only samples with 2 out of 2 positive RT-qPCR reactions are presented. Data are shown as the mean of the calculated N gene copy number per ml of liquid sample or per mg of a 20% tissue homogenate.

### Histopathology

Tissue samples from the respiratory tract (nasal cavity [rostral, middle and deep turbinates following decalcification with Immunocal™ Decalcifier [StatLab, McKinney, TX] for 4-7 days at room temperature]), trachea, and lungs as well as various other extrapulmonary tissues (liver, spleen, kidneys, heart, pancreas, gastrointestinal tract [stomach, small intestine including Peyer’s patches and colon], cerebrum [including olfactory bulb], tonsils and numerous lymph nodes were routinely processed and embedded in paraffin. Four-micron tissue sections were stained with hematoxylin and eosin following standard procedures. Multiple independent veterinary pathologists (blinded to the treatment groups) examined the slides and morphological descriptions were provided.

### SARS-CoV-2-specific RNAscope® in situ hybridization (RNAscope® ISH)

RNAscope® ISH was performed as previously described (12) using an anti-sense probe targeting the spike protein gene (S; nucleotide sequence: 21,563-25,384) of SARS-CoV-2, USA-WA1/2020 isolate (GenBank accession number MN985325.1) which was designed (Advanced Cell Diagnostics [ACD], Newark, CA, USA) and used as previously described (18). Four-micron sections of formalin-fixed paraffin-embedded tissues were subjected to *in situ* hybridization. Lung sections from a SARS-CoV-2-infected hamster were used as positive assay controls.

### SARS-CoV-2-specific immunohistochemistry (IHC)

IHC was performed as previously described (12) on four-micron sections of formalin-fixed paraffin-embedded tissue mounted on positively charged Superfrost® Plus slides and subjected to IHC using a SARS-CoV-2-specific anti-nucleocapsid rabbit polyclonal antibody (3A, developed by our laboratory) with the method previously described (18). Lung sections from a SARS-CoV-2-infected hamster were used as positive assay controls.

#### Virus neutralizing antibodies

Virus neutralizing antibodies in sera were determined using microneutralization assay as previously described (12). Briefly, heat inactivated serum samples were subjected to 2-fold serial dilutions starting at 1:20 in duplicate. Then, 100 TCID_50_ of SARS-CoV-2 virus in DMEM culture media was added 1:1 to the sera dilutions and incubated for 1 h at 37 ^°^C, then cultured on Vero E6 cells in 96-well plates. The corresponding SARS-CoV-2-negative cat sera, virus only and media only controls were also included in the assay. The neutralizing antibody titer was recorded as the highest serum dilution at which at least 50% of wells showed virus neutralization based on the appearance of CPE observed under a microscope at 72 h post infection.

#### Detection of SARS-CoV-2 antibodies by indirect ELISA

To detect SARS-CoV-2 antibodies in sera, indirect ELISAs were performed with the recombinant viral proteins, nucleoprotein (N) and the receptor-binding domain (RBD), as previously described (12). Briefly, wells were coated with 100 ng of the respective recombinant protein and serum samples were pre-diluted 1:400 for the assay. The cut-off for a sample being called positive was determined as follows: Average OD of negative serum + 3X standard deviation. Everything above this cut-off was considered positive.

## Results

### 1. Primary infected cats shed SARS-CoV-2, develop histologic lesions in the respiratory tract, and seroconvert while remaining clinically asymptomatic

Two independent studies were conducted by inoculating 3 and 5 principal cats with SARS-CoV-2 in the first and second studies, respectively, and introducing 2 sentinel contact cats at 1 DP2C in the second study (12; **Figure 1**). Similar to our previous study (12), primary SARS-CoV-2 challenged cats shed virus from nasal, oral and rectal cavities and efficiently transmitted the virus to the naïve sentinels (**Figure 2a-c**). Viral RNA was found in multiple tissues of the 2 principal infected cats euthanized at 4 DPC, including the URT, LRT, GIT, lymphatic organs, heart and olfactory bulb (**Figure 3**). Histologic lesions were restricted to the upper respiratory tract and bronchial tree as previously described (12). Within the nasal cavity, there was evidence of neutrophilic rhinitis of variable intensity with numerous neutrophils infiltrating the lamina propria, transmigrating through the lining epithelium, and accumulating in the lumen along with cellular debris (**Figure 4**). The respiratory epithelium showed occasional sloughing (erosion) and attenuation. Seromucinous nasal glands were largely spared, with only rare glands within inflamed areas containing few luminal neutrophils or cell debris. SARS-CoV-2 antigen and viral RNA were multifocally detected within individualized or clusters of squamous (rostral turbinates) and respiratory epithelial cells (intermediate and deep turbinates) via IHC and ISH. Even though the olfactory neuroepithelium was histologically unremarkable, viral antigen and RNA were segmentally detected (**Figure 4**). Lesions in the trachea and bronchial tree were characterized by multifocal lymphohistiocytic and neutrophilic tracheobronchoadenitis with necrosis of the glandular epithelium and intralesional SARS-CoV-2 antigen and RNA as previously described (**Figure 4**; 12). At 21 DPC, viral shedding had subsided in both principal and contact primary infected animals, except for a single positive oropharyngeal swab detected from one of the principal infected cats (**Figure 2b**). Limited viral RNA was detected in tissues of the URT, lymphatic organs and olfactory bulb, but not in the nasal wash, BALF, lung or digestive tract tissues of the principal infected cat euthanized at 21 DPC (**Figure 3**). All cats including principal and sentinel primary infected animals had detectable virus-specific and virus neutralizing antibodies by 21 DPC/0 DP2C (**Tables 2 and 3**). At this timepoint, the intense neutrophilic rhinitis noted at 4 DPC was replaced by multifocal lymphoid aggregates in the lamina propria with only rare and localized areas of minimal neutrophilic inflammation. The respiratory mucosa and olfactory neuroepithelium were unremarkable and no viral antigen or RNA were detected within the nasal cavity at this timepoint (**Figure 4**). In the trachea and bronchi, there was no evidence of damage to tracheal and bronchial glands as noted at 4 DPC, with no viral antigen or RNA detected. Tracheal glands were only separated by mild numbers of lymphocytes and plasma cells, and peribronchial lymphoid aggregates were noted along the bronchial tree (**Figure 4**). Finally, no lesions were identified in extrapulmonary tissues other than lymphoid hyperplasia within lymphoid organs.

**Table 2.** Re-infection study 1. SARS-CoV-2 specific and virus neutralizing antibody responses following re-challenge.

| Anti-N antibodies |  |  |  |  | Anti-RBD antibodies |  |  |  | Virus neutralizing antibodies |  |  |  |
| --- | --- | --- | --- | --- | --- | --- | --- | --- | --- | --- | --- | --- |
| DP2C | 0 | 1 | 3 | 4 | 0 | 1 | 3 | 4 | 0 | 1 | 3 | 4 |
| *Re-challenged principal infected cats* |  |  |  |  |  |  |  |  |  |  |  |  |
| 328 | na | + | + | + | na | + | + | + | 160 | 80 | 160 | 160 |
| *Re-challenged sentinels* |  |  |  |  |  |  |  |  |  |  |  |  |
| 272 | + | + | + | + | + | + | + | + | 160 | 80 | 160 | 80 |
| 903 | + | + | + | + | + | + | + | + | 160 | 160 | 80 | 160 |

DP2C=days post second challenge, which occurred 21 days after primary challenge; na=not available; N=nucleoprotein; RBD=receptor binding domain; +/-=OD close to cut-off

**Table 3.**
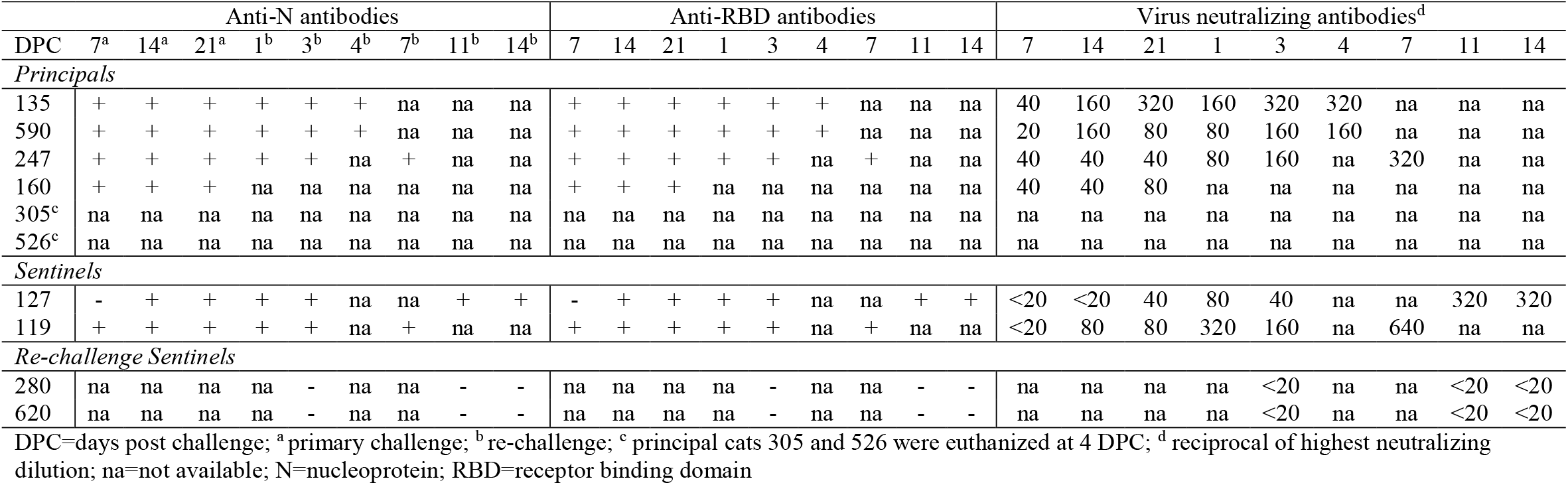
Re-infection study 2. SARS-CoV-2 specific and neutralizing antibody responses following primary challenge and re-challenge.

**Figure 1.**
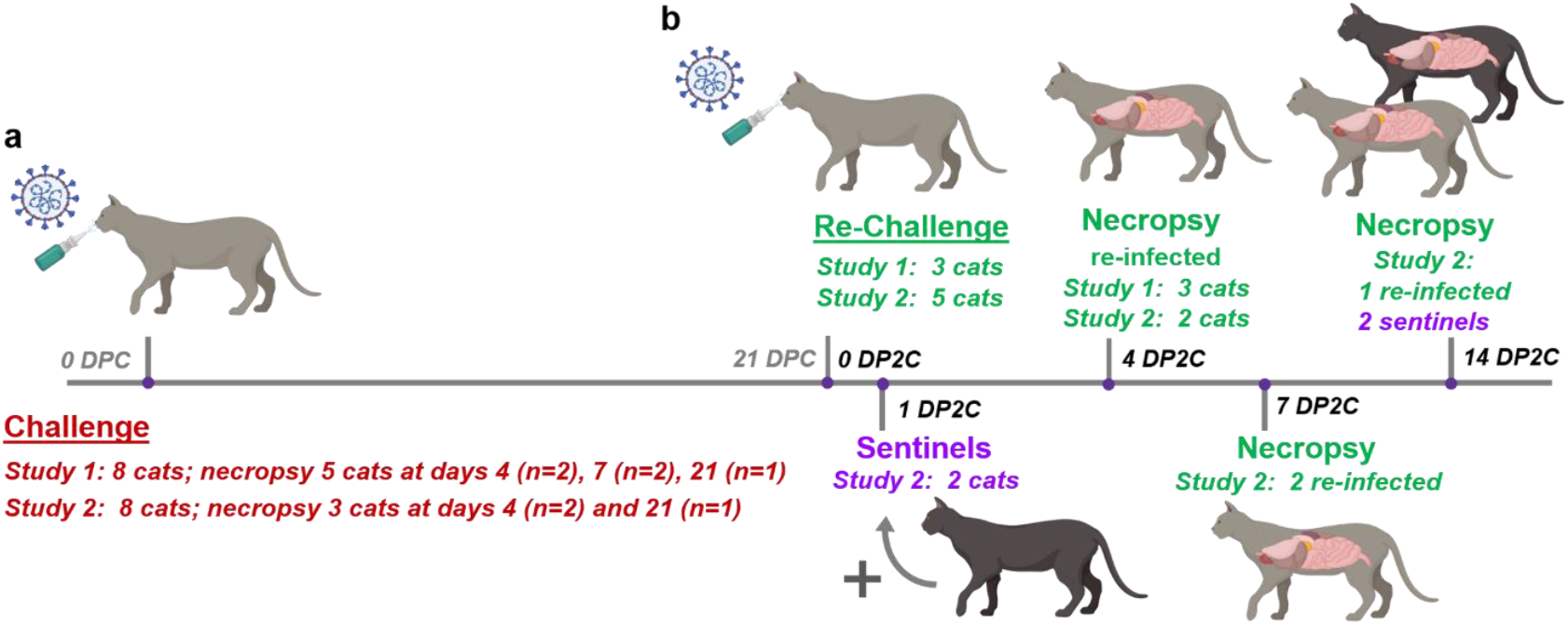
Re-infection study design. (a) In each of 2 studies, 6 cats were inoculated with SARS-CoV-2 and 2 sentinel contact cats introduced the following day post primary challenge (DPC). Necropsy was performed on principal infected cats at 4, 7 and 21 DPC. (b) At 21 DPC, cats were re-challenged with SARS-CoV-2 at the same dose as primary challenge, and 2 sentinels introduced at 1 day post second challenge (DP2C). Necropsy of principal re-infected cats were performed at 4, 7 and 14 DP2C, and the sentinels at 14 PD2C.

**Figure 2.**
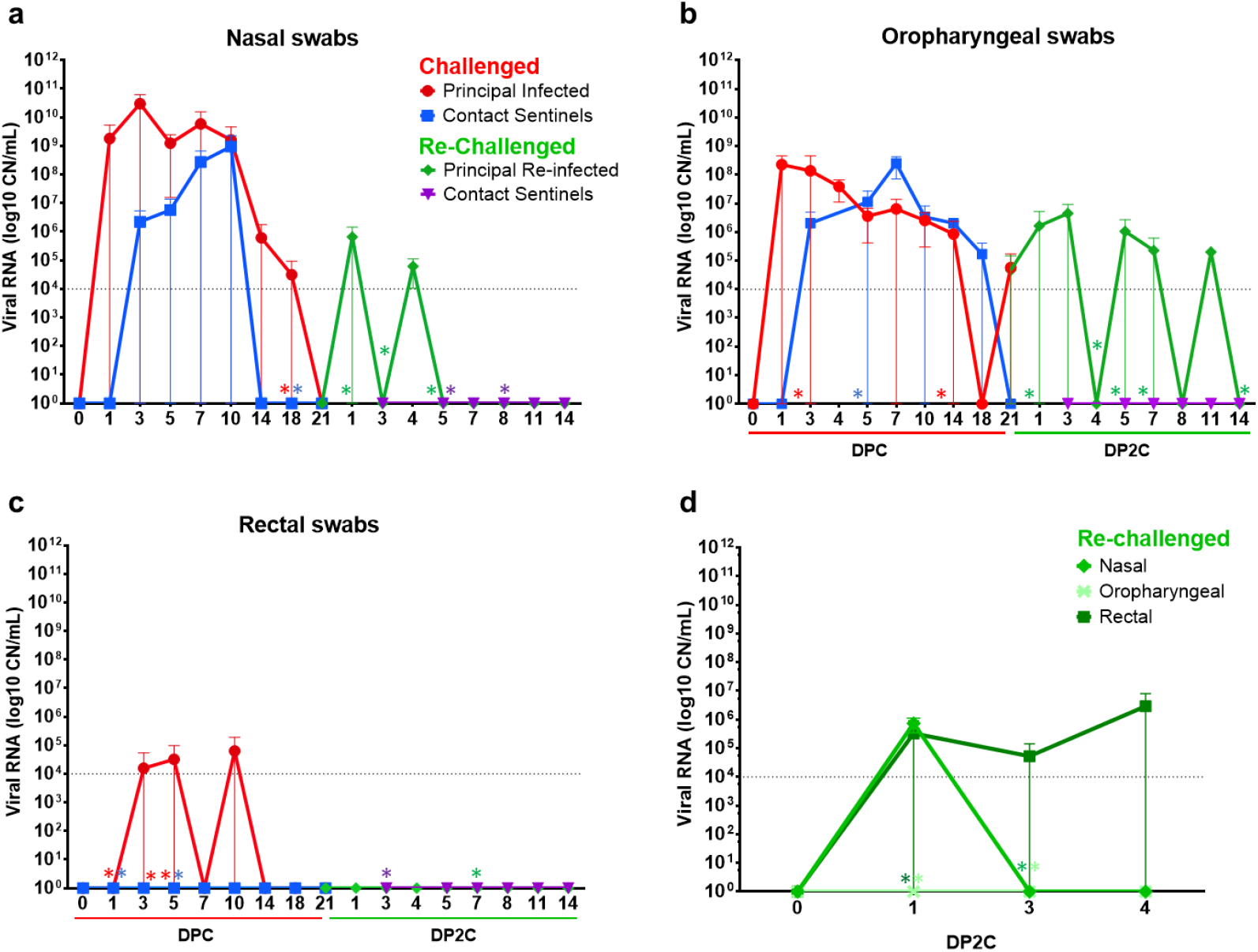
Viral shedding from SARS-CoV-2 infected and re-infected cats. RT-qPCR was performed on nasal (a), oropharyngeal (b) and rectal (c) swabs collected from cats at the indicated days following primary challenge (DPC) and re-challenge (DP2C) from the second re-infection study. (d) Nasal, oropharyngeal and rectal swabs collected from re-challenged cats from the first re-infection study. Mean and standard deviation of viral RNA copy number (CN) per mL based on the nucleocapsid gene are shown. Asterisks (*) indicate samples with 1 out of 2 of the RT-qPCR reactions below the limit of detection, indicated by the dotted line.

**Figure 3.**
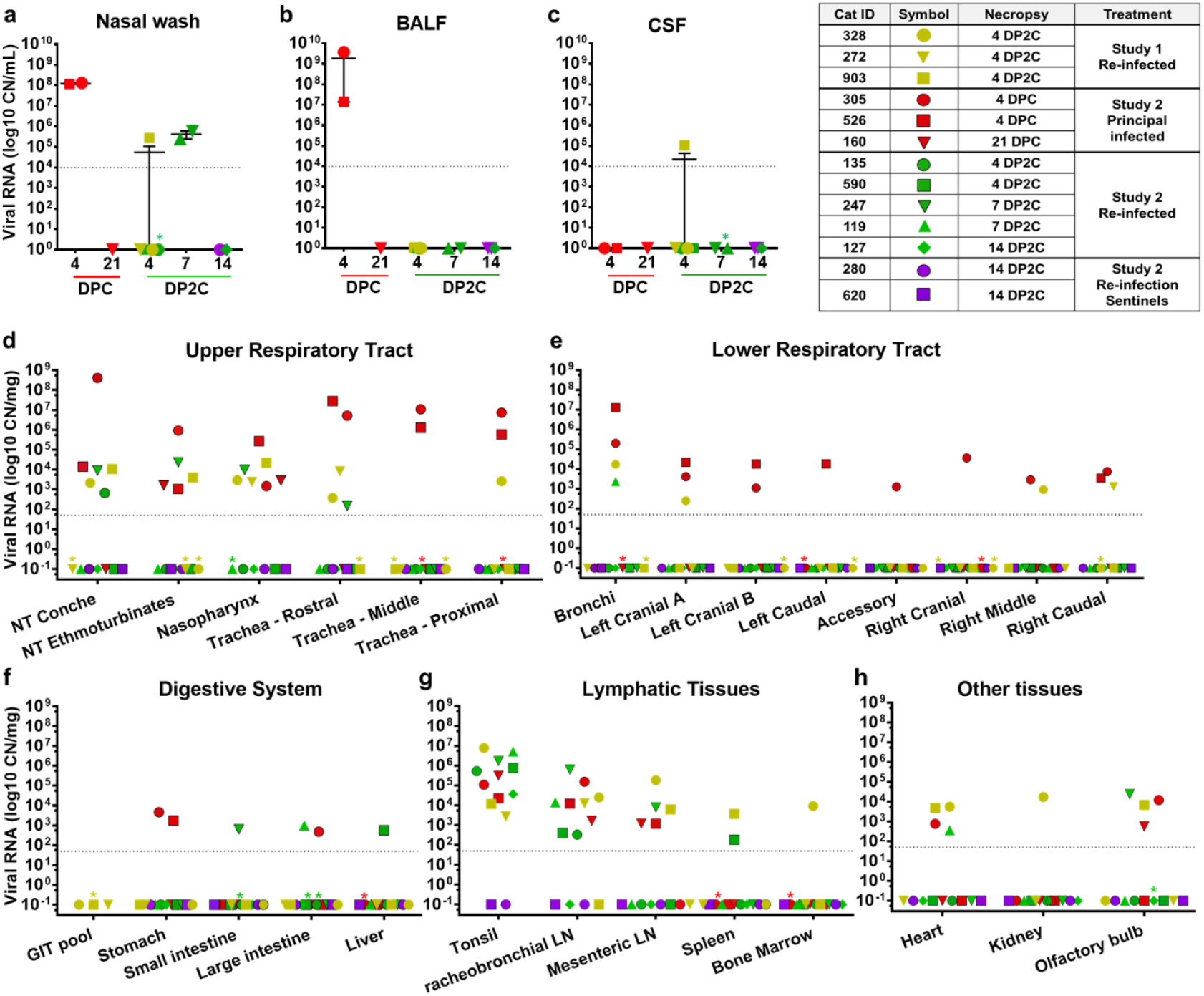
SARS-CoV-2 RNA detected in various tissues from infected and re-infected cats. RT-qPCR was used to detect the presence of SARS-CoV-2 in various tissues of cats euthanized at the indicated days after primary challenge (DPC) and re-challenge (DP2C). Viral RNA copy number (CN) per mL (a-c) or mg (d-h) based on the nucleocapsid gene are plotted for individual animals. Colored symbols corresponding to cat ID numbers, day of necropsy and study are indicated in the figure key. Asterisks (*) indicate samples with 1 out of 2 of the RT-qPCR reactions below the limit of detection, indicated by the dotted line.

**Figure 4.**
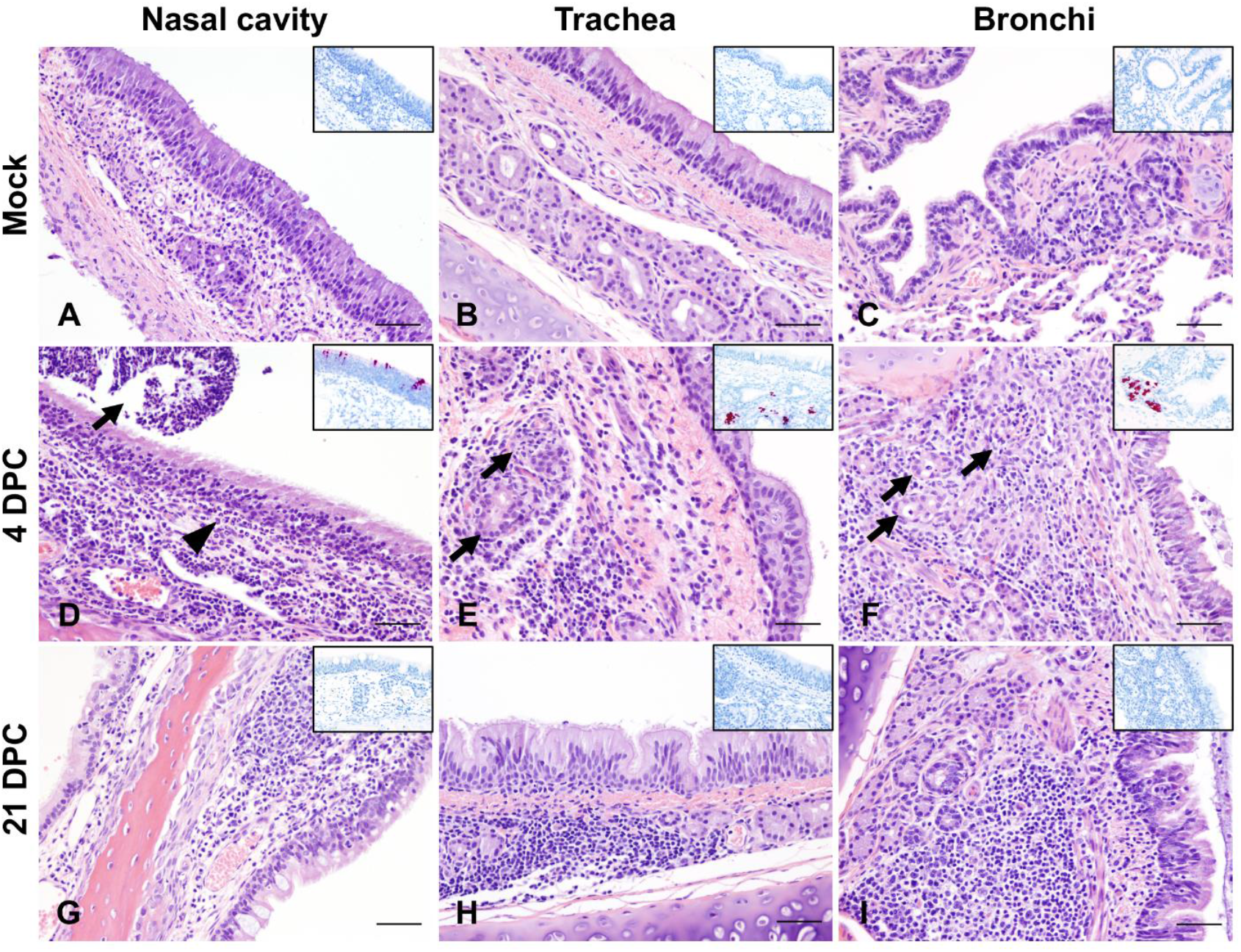
Histologic lesions and SARS-CoV-2 distribution as determined by *in situ* hybridization in the respiratory tract of primary infected cats. Mock-infected (A-C), 4 (D-F), and 21 days post-challenge (DPC; G-I) with SARS-CoV-2. At 4 DPC, intense neutrophilic rhinitis with luminal exudate (D, arrowhead and arrow, respectively) and lymphohistiocytic and neutrophilic tracheobronchoadenitis with necrosis and obliteration of seromucinous glands (E and F, arrows) were characteristic. Viral antigen and RNA (red) localized within the nasal respiratory epithelium, olfactory neuroepithelium (D inset), and affected tracheal and bronchial glands (E and F insets). At 21 DPC, histologic changes were limited to lymphoid aggregates in the lamina propria of the nasal turbinates (G), trachea (H) and bronchi (I), and no viral antigen or RNA was detected (insets). H&E and Fast Red, 200X total magnification. Bar = 100 μm.

### 2. Re-infection of cats with SARS-CoV-2 leads to asymptomatic and limited viral shedding

At 21 DPC/0 DP2C, 3 cats from the first study and 5 cats of the second study were re-challenged with SARS-CoV-2 (**Figure 1**). In the first study, viral shedding was detected from nasal swabs at 1 DP2C and from rectal swabs at 1 to 4 DP2C, but not from oropharyngeal swabs of the 3 re-challenged cats (**Figure 2d**). In contrast, re-challenged cats from the second study shed viral RNA from the nasal cavity at 1 through 4 DP2C and from the oropharyngeal cavity at 1 through 11 DP2C; however, no viral RNA was detected from rectal swabs after re-challenge in study 2 (**Figure 2a-c**).

In addition, viral RNA was detected in the nasal wash, URT and GIT tissues of re-infected cats euthanized at 4 and 7 DP2C (**Figure 3a**,**d**,**f**). Limited viral RNA was detected in the LRT of some animals and some LRT tissues of cats euthanized after re-infection at 4 and 7 DP2C, including the bronchi and 3 of the 7 lung sections collected (**Figure 3b**,**e**). Viral RNA was also found in the lymphatic tissues, heart and olfactory bulb of the re-infected cats at 4 and 7 DP2C at similar levels detected in the cats during primary infection (**Figure 3**). CSF from one of the re-challenged cats euthanized at 4 DP2C was also positive (**Figure 3c**). By 14 DP2C, viral RNA was only detected in the tonsil of the remaining principal re-challenged cat. No viral RNA was detected from the blood of primary SARS-CoV-2 challenged or re-challenged cats (data not shown).

No significant clinical symptoms including fever or weight changes were observed for any of the primary SARS-CoV-2 challenged or re-challenged cats during the course of these studies. On the day of re-challenge, all cats had virus neutralizing antibodies with titers ranging from 1:40 to 1:320 (**Tables 2 and 3**). A marked increase in neutralizing antibody titers was observed at 7 to 11 DP2C, suggestive of an anamnestic immune response after re-infection with SARS-CoV-2.

Histologic changes within the upper and lower respiratory tracts, as well as extrapulmonary tissues were evaluated in re-infected cats. While there was evidence of minimal to mild neutrophilic rhinitis (particularly on rostral and intermediate turbinates) at 4 DP2C, this was not as intense as that observed at 4 DPC. In all re-infected cats (4, 7 and 14 DP2C), multiple lymphoid aggregates were noted in the nasal passages similar to those noted at 21 DPC (**Figure 5**). No alterations within the respiratory mucosa (other than occasional transmigrating lymphocytes) and olfactory neuroepithelium were noted in nearly all re-infected animals. In a single cat (#590-4 DP2C), few neutrophils infiltrated a localized area of the olfactory neuroepithelium. No viral antigen or RNA were detected in the nasal cavity at any timepoint following re-infection (**Figure 5**). In the trachea and bronchi of re-infected cats, there was no evidence of alterations to tracheal and bronchial glands as noted at 4 DPC, with no viral antigen or RNA detected. Mild numbers of lymphocytes and plasma cells infiltrated between tracheal and bronchial glands, and frequently formed lymphoid aggregates along the bronchial tree, interpreted as hyperplastic bronchial-associated lymphoid tissue (BALT, **Figure 5**).

**Figure 5.**
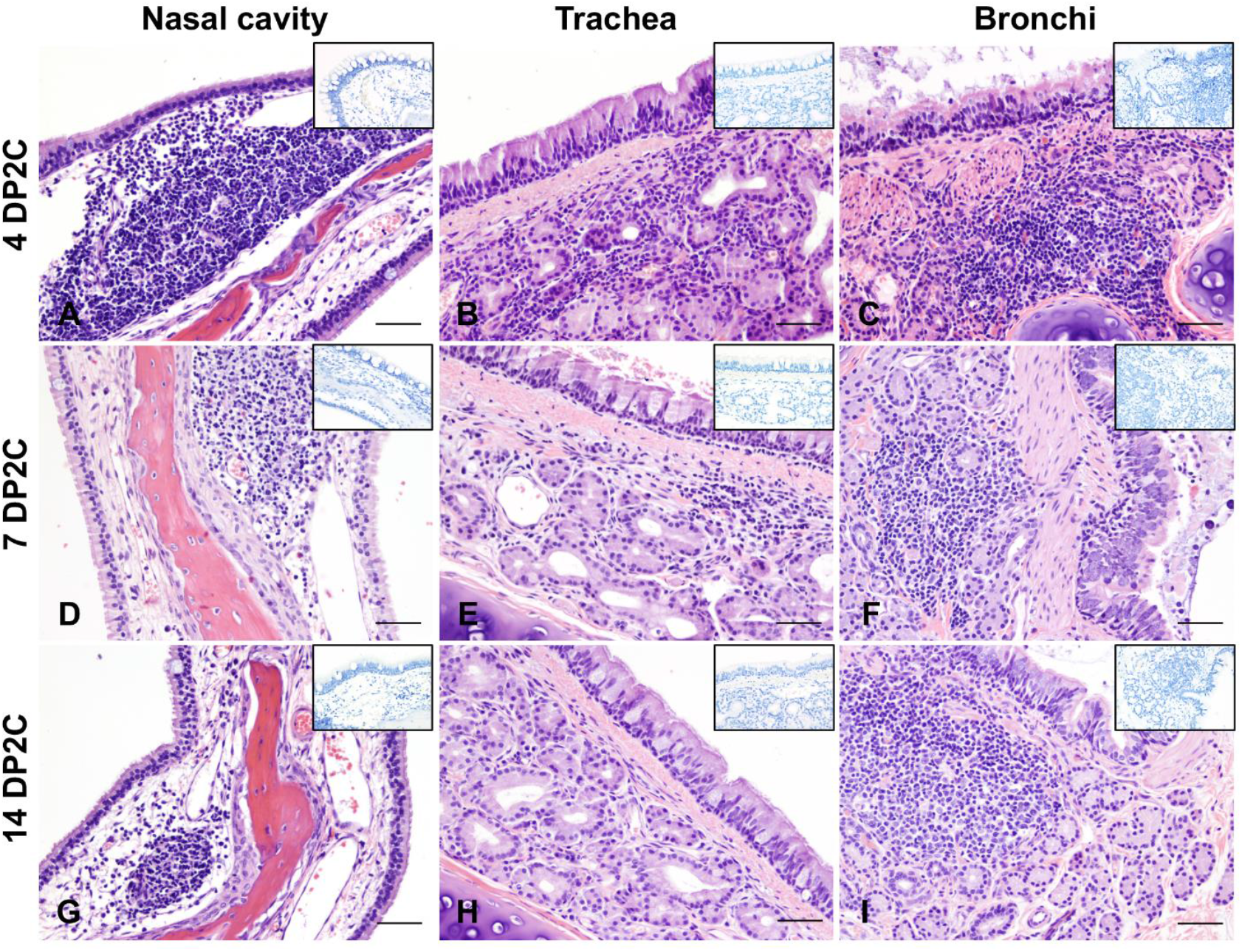
Histologic lesions and SARS-CoV-2 distribution as determined by *in situ* hybridization in the respiratory tract of re-challenged cats. 4 (A-C), 7 (D-F), and 14 days post-second challenge (DP2C; G-I) with sARS-CoV-2. At all timepoints, variable lymphoid aggregates expanded the lamina propria of the nasal turbinates (A, D, G), trachea (B, E, H) and bronchi (C, F, I), and no viral antigen or RNA was detected (insets). H&E and Fast Red, 200X total magnification. Bar = 100 μm.

While no significant histologic alterations were evident in the pulmonary parenchyma of primary infected cats and most of the re-infected cats (5/8), a localized area of reparative alveolitis was identified in a single bronchopulmonary segment in re-infected cats #590 (4 DP2C), #119 (7 DP2C) and #127 (14 DP2C). The affected area was characterized by early (#590) to florid (#119 and #127) alveolar type 2 cell hyperplasia with few infiltrating mononuclear cells (lymphocytes and histiocytes) and small amounts of sloughed cell debris. No SARS-CoV-2 antigen or RNA were detected in the affected area or elsewhere in the pulmonary parenchyma (data not shown). No histologic changes in extrapulmonary organs were observed other than lymphoid hyperplasia within lymphoid organs.

### 3. Re-infected cats do not transmit SARS-CoV-2 to sentinels

In the second re-infection study, 2 sentinel SARS-CoV-2 antibody-negative contact cats were introduced and co-mingled with the re-challenged cats at 1 DP2C in order to determine if virus transmission occurred following re-infection. The sentinels introduced after re-infection had temperatures slightly above 39^°^C at 2 to 4 DP2C but were otherwise within normal range and remained asymptomatic up to 14 DP2C (**supplemental figure 1**). No viral shedding from nasal, oropharyngeal and rectal cavities was observed in these animals (**Figure 2a-c**) and no viral RNA was detected in tissues from either of the sentinel cats examined at 14 DP2C (**Figure 3**). Furthermore, the naïve sentinel cats co-housed with the re-infected cats for the 13 days after re-challenge did not seroconvert (**Tables 2 and 3**). Histologic evaluation of the nasal cavity identified multiple lymphoid aggregates similar to those noted in cats at 21 DPC and following re-infection with SARS-CoV-2. Minimal infiltrating lymphocytes and plasma cells along with sporadic lymphoid aggregates were noted in the submucosa of the trachea and along the bronchial tree, with no histologic changes in the pulmonary parenchyma. Viral antigen and RNA were not detected in tissues from sentinel cats.

## Discussion

Knowledge of SARS-CoV-2 viral evolution, immunology and the longevity of immune protection against re-infection is still limited. Cases of SARS-CoV-2 re-infections in humans have been recently reported (19-25). Although the reported cases are relatively few, the true percentage and frequency of SARS-CoV-2 re-infections remain unclear. Furthermore, the outcome of a re-infection in terms of clinical disease and transmission is still not clearly understood. While identifying and characterizing re-infection in patients naturally infected or vaccinated and re-exposed to SARS-CoV-2 remains important, studies using susceptible animal models allows for more systematic, in-depth investigation which can provide insight into SARS-CoV-2 re-infections in humans. Therefore, we performed an in-depth investigation of experimental re-infection in domestic cats as well as transmission following re-infection with SARS-CoV-2.

Our results indicate that a primary SARS-CoV-2 infection was mostly resolved within 21 DPC. At 21 DPC, viral RNA shedding had subsided in the nasal and rectal cavities (**Figure 2**), no viral RNA or antigen were detected in the upper and lower respiratory tract with the exception of nasal turbinate and nasopharynx homogenates (**Figure 3 and 4**), and the histologic changes seen at early stages of infection (i.e., 4 and 7 DPC), characterized by intense neutrophilic rhinitis and tracheobronchoadenitis with necrosis of seromucinous glands of the trachea and bronchi, had completely resolved with only occasional lymphoid aggregates along segmental bronchi. Following re-challenge, our results indicate that cats were at least partially protected from re-infection. All cats had circulating neutralizing antibodies at the time of re-challenge (**Tables 2 and 3**), and viral RNA was detected from fewer tissues and at lower levels in the re-infected cats compared to primary infected cats examined at the same day after virus challenge (12; **Figures 2 and 3**). In our first re-infection study, viral RNA was detected in nasal washes, the URT and some LRT tissues of the 3 re-infected cats examined at 4 DP2C, suggesting that limited infection mainly in the URT might occur after SARS-CoV-2 re-challenge. Interestingly, in our second study, nasal washes, URT and GIT tissues of the 2 cats examined at 7 DP2C but not at 4 DP2C were RNA positive; this might suggest that the viral RNA detected in the URT and GIT may not be residual viral RNA from the SARS-CoV-2 challenge but replicating virus following re-challenge. Limited viral shedding was also detected from nasal and rectal cavities up to 4 DP2C and oropharyngeal swabs up to 11 DP2C. However, no viable virus was isolated from the swab samples of the re-infected cats, and the 2 sentinel contact cats co-housed with the re-challenged cats for 13 days remained SARS-CoV-2 negative and did not seroconvert. Interestingly, re-infected cats did not develop intense neutrophilic rhinitis or intense, acute inflammation and necrosis targeting tracheobronchial glands when compared to primary infected cats at 4 DPC, and no viral antigen and RNA was detected within respiratory tract tissues following re-infection. The lymphoid aggregates frequently noted within the nasal passages and throughout the bronchial tree of re-infected cats resemble hyperplastic nasal and bronchial-associated lymphoid tissue (NALT and BALT), most likely a non-specific change following exposure to airborne pathogens. Whether this change represents a specific host immune response that either remained following the initial exposure to SARS-CoV-2 and expanded following re-exposure is unclear, since similar lymphoid clusters, albeit at lower frequency, were noted in the sentinel cats and in one of the mock-infected cats. A larger sample size would be necessary in order to confidently determine the frequency and distribution of this histopathological finding in outbred cats as those used in this study.

An additional and unexpected histologic lesion when compared to primary infected cats corresponded to the focal reparative alveolitis in three of the re-challenged cats. Unfortunately, the association of this histologic change with SARS-CoV-2 re-infection cannot be unequivocally confirmed. However, its association with SARS-CoV-2 infection is unlikely due to several reasons: 1) SARS-CoV-2 does not show a specific tropism for feline bronchial/bronchiolar or alveolar epithelial cells (12); 2) no viral antigen or viral RNA (both via ISH and RT-qPCR) were detected within these lesions; 3) histologic changes suggestive of alveolar injury and repair were regional and not generalized and were not associated with clinical signs of disease; 4) the alveolar lesions were not present in all re-challenged cats. Other possible causes such as immune-mediated responses, or other inhaled antigens from the inoculum or environment should be considered. Additional experiments with a larger sample size are warranted to further investigate the mechanisms of immune protection from SARS-CoV-2 and the development of pulmonary immunity in cats. Taken together, these results indicate that experimental re-infection of cats with SARS-CoV-2 results in partial protection from infection with limited infection primarily of the URT and GIT that was resolved by 14 DP2C; regardless, re-challenged animals did not shed viable virus at sufficient levels for efficient transmission to sentinels.

Other SARS-CoV-2 re-infection studies in cats, ferrets and rhesus macaques also showed at least partial to full protection from re-exposure at 4 weeks after primary challenge (26-28). In a study by Bosco-Lauth and colleagues (2020), 3 adult cats 5-8 years of age were experimentally infected with SARS-CoV-2 and re-challenged 28 days later. In that study, no nasal or oral shedding of virus was observed during the 7 days after re-challenge and it was concluded that the animals were resistant to SARS-CoV-2 re-infection (15). In our studies, we also did not detect live virus but did detect viral RNA shedding. Similar to Bosco-Lauth et al., we also observed a marked increase in neutralizing antibodies following SARS-CoV-2 re-challenge, suggestive of an anamnestic immune response. Similar results to ours were obtained in re-infection studies with non-human primates (NHPs) by two different groups (27, 28). After the initial SARS-CoV-2 clearance, animals were re-challenged with SARS-CoV-2 and showed significant reductions in viral loads in bronchoalveolar lavage and nasal mucosa as compared to the primary infection (27, 28). Anamnestic immune responses after re-challenge suggested that protection was mediated by immunologic control. The authors concluded that SARS-CoV-2 infection in rhesus macaques led to SARS-CoV-2 specific immune responses and provided protection against re-challenge. The residual levels of subgenomic mRNA in nasal swabs and anamnestic immune responses after SARS-CoV-2 re-challenge suggest that protection was mediated by immunologic control but was not sterilizing (27). While these studies indicate that SARS-CoV-2 infection induces immune responses that provide at least some protection against re-infection in the short-term, the longevity of immune protection and the extent of cross-protection against divergent SARS-CoV-2 isolates that emerge remain unclear.

Some insight into the longevity of immune responses and protection from re-infections might be derived from studies on similar coronaviruses. Studies on recovered patients infected with Severe Acute Respiratory Syndrome coronavirus (SARS-CoV) and Middle East Respiratory Syndrome coronavirus (MERS-CoV) indicate that virus-specific and neutralizing antibodies can persist for up to 2 to 3 years before significantly declining; still, protection against re-infection in such seropositive individuals is not known (29-35). MERS-CoV re-infections in seropositive camels has been shown to occur (36, 37). A 35 year-long study investigating re-infections in individuals with seasonal human coronaviruses that typically result in minor disease found re-infections occurred frequently around a year after the previous infection indicating immunity is rather short-lived (38). Further studies are needed to understand the long-term kinetics of immune responses to SARS-CoV-2 and how these responses relate to protection from re-infection, and in cases of only partial protection, the risk of transmission and developing clinical disease. This work is also critical for determining the risk of companion animals as reservoirs for perpetuating spread of SARS-CoV-2 and may also serve as a model to study SARS-CoV-2 immunity and re-infection in humans.

## Conclusions

Here, we demonstrate that experimental SARS-CoV-2 infection in cats induces a protective immune response providing partial non-sterilizing immune protection from re-infection. Furthermore, we show that re-infected cats did not shed virus at sufficient levels to infect co-housed, naïve animals. Future studies are aimed at understanding the longevity of immune protection against SARS-CoV-2. Nonetheless, these results suggest that immunological approaches to prevent and potentially treat SARS-CoV-2 are possible.

## Acknowledgments

We thank the staff of KSU Biosecurity Research Institute, the histological laboratory at the Kansas State Veterinary Diagnostic Laboratory (KSVDL), the CMG staff and Sabarish Indran, Gleyder Roman-Sosa, Yonghai Li, and Emily Gilbert-Esparza at KSU. We also thank the staff at the Histology and Immunohistochemistry section of the Louisiana Animal Disease Diagnostic Laboratory (LADDL) for assistance with this study. The SARS-CoV-2 strain USA-WA1/2020 was obtained through BEI Resources (catalogue # NR-52281).

Mention of trade names or commercial products in this publication is solely for the purpose of providing specific information and does not imply recommendation or endorsement by the U.S. Department of Agriculture. USDA is an equal opportunity provider and employer.

## Declaration of conflict of interest

The authors declared no potential conflicts of interest with respect to the research, authorship, and/or publication of this article

## Funding

Funding for this study was provided through grants from NBAF Transition Funds from the State of Kansas, the NIAID Centers of Excellence for Influenza Research and Surveillance under contract number HHSN 272201400006C, and the Department of Homeland Security Center of Excellence for Emerging and Zoonotic Animal Diseases under grant number HSHQDC 16-A-B0006 to JAR. This study was also partially supported by the Louisiana State University, School of Veterinary Medicine start-up fund (PG 002165) to UBRB and the U.S. Department of Agriculture, Agricultural Research Service (58-32000-009-00D) to WCW, by the Center for Research for Influenza Pathogenesis (CRIP), a NIAID supported Center of Excellence for Influenza Research and Surveillance (CEIRS, contract # HHSN272201400008C), and by the generous support of the JPB Foundation, the Open Philanthropy Project (research grant 2020-215611 [5384]) and anonymous donors to AG-S.

---

**Supplementary figure 1.**
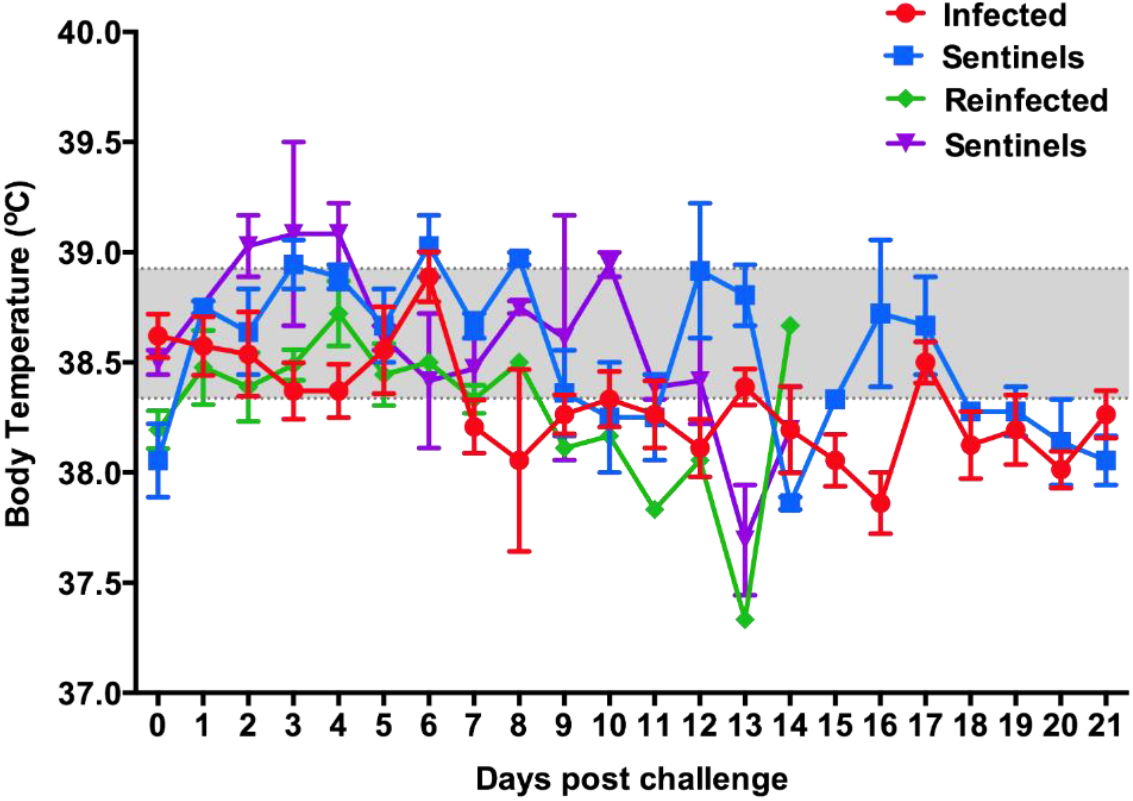

## Notes

### Competing Interest Statement

The authors have declared no competing interest.

